# A mitochondrial iron-sensing pathway regulated by DELE1

**DOI:** 10.1101/2022.04.14.488327

**Authors:** Yusuke Sekine, Ryan Houston, Evelyn Fessler, Lucas T Jae, Derek P Narendra, Shiori Sekine

## Abstract

The heme-regulated kinase HRI is activated under heme/iron deficient conditions; however, the underlying molecular mechanism is incompletely understood. Here, we show that iron deficiency-induced HRI activation involves a heme-independent mechanism that requires the mitochondrial protein DELE1. Notably, mitochondrial import of DELE1 and its subsequent protein stability are regulated by iron availability. Under steady state conditions, DELE1 is degraded by the mitochondrial matrix-resident protease LONP1 soon after mitochondrial import. Upon iron chelation, DELE1 import is arrested, thereby stabilizing DELE1 on the mitochondrial surface to activate the HRI-mediated integrated stress response (ISR). Moreover, depletion of the mitochondrial ABC transporter ABCB7 that is involved in iron-sulfur cluster (ISC) metabolism markedly abrogates iron deficiency-induced ISR activation, suggesting the possible involvement of ISC-related molecules in this activation. Our findings highlight mitochondrial import regulation of DELE1 as the core component of a previously unrecognized iron monitoring system that connects the mitochondria to the cytosol.

## Introduction

The integrated stress response (ISR) is a highly conserved program that plays a central role in the cellular adaptation to stress conditions (Costa-Mattioli and Walter, 2020; Pakos-Zebrucka et al., 2016). In mammalian cells, this response is initiated by four stress-responsive kinases including HRI, PKR, PERK, and GCN2, which are activated by different stresses (Donnelly et al., 2013; Houston et al., 2020). These kinases phosphorylate a single substrate, the α subunit of the translation initiation factor eIF2 (eIF2α), thereby attenuating global mRNA translation, while activating the transcription factor ATF4, a master regulator of the transcriptional response for the ISR.

The eIF2α kinase HRI is expressed predominantly in erythroid cells, governing the ISR during terminal erythropoiesis (Chen and Zhang, 2019). Early studies using rabbit reticulocyte lysates revealed that HRI is a major suppressor of protein synthesis, and that the addition of hemin (heme) to the lysates reverses HRI-mediated translational inhibition (Bruns and London, 1965; Ranu et al., 1976; Ranu and London, 1976; Trachsel et al., 1978). Subsequent *in vitro* studies using purified HRI demonstrated that the direct binding of heme on HRI modulated the conformation of HRI thereby suppressing its kinase activity (Fagard and London, 1981; Igarashi et al., 2008; Miksanova et al., 2006; Rafie-Kolpin et al., 2000). These findings led to a current model of HRI regulation in which HRI is activated through dissociation of the inhibitory heme from HRI when intracellular levels of heme decline. It has been shown that HRI knockout (KO) mice do not exhibit significant erythroid abnormalities under standard dietary condition (Han et al., 2001). However, when challenged with an iron deficient diet, these mice develop a characteristic anemia with globin aggregation in red blood cells (Han et al., 2001). One potential explanation for these iron-dependent *in vivo* effects is that iron deficiency concomitantly decreases heme levels, and this fall in heme is the mechanism of HRI activation. However, such conjectures have remained unproven.

It has been shown that various types of mitochondrial stress, including membrane potential loss, inhibition of OXPHOS, and perturbation of mitochondrial proteostasis, activate the ISR (Munch and Harper, 2016; Quiros et al., 2017; Taniuchi et al., 2016). Recent cell genetic screens have demonstrated that HRI is required for the mitochondrial stress-triggered ISR activation (Fessler et al., 2020; Guo et al., 2020). Moreover, the mitochondria resident proteins DELE1 and OMA1 have been identified as upstream regulators of HRI (Fessler et al., 2020; Guo et al., 2020). These studies have demonstrated that upon treatment with various mitochondrial stressors (including agents that cause mitochondrial membrane depolarization, and inhibitors of mitochondrial complex V or the mitochondrial matrix chaperon), the mitochondrial protease OMA1 cleaves DELE1, and the cleaved form of DELE1 is released into the cytosol where it interacts with HRI, activating the ISR. Thus, the OMA1-DELE1-HRI pathway constitutes a retrograde signal from mitochondria to the cytosolic ISR under mitochondrial stress conditions. However, the functional link between this newly identified mitochondria-dependent pathway and the classic heme-dependent HRI regulation model remains unclear.

Accumulating evidence indicates that mitochondrial import of certain proteins is precisely regulated in response to mitochondrial stress. Importantly, this regulation is often used to signal mitochondrial stress to other subcellular compartments such as the cytosol and the nucleus. A well characterized example is the Parkinson’s disease (PD)-linked mitochondrial kinase PINK1, whose import arrest and stabilization on the outer mitochondrial membrane (OMM) upon mitochondrial depolarization triggers Parkin-mediated mitophagy (Matsuda et al., 2010; Narendra et al., 2010; Sekine and Youle, 2018). Furthermore, it has been shown in *C. elegans* that suppression of mitochondrial import of the transcription factor ATFS-1 by proteotoxic stress leads to the translocation of ATFS-1 to the nucleus, which is central for the transcriptional response known as the mitochondrial unfolded protein response (mtUPR) (Anderson and Haynes, 2020). Mitochondrial import arrest of ATFS-1 under proteotoxic stress conditions requires the peptide export activity of HAF-1, an inner mitochondrial membrane (IMM)-resident ATP-binding cassette (ABC) transporter (Haynes et al., 2010; Nargund et al., 2012). Peptides are thought to be derived from unfolded proteins digested by mitochondrial protease CLPP (Haynes et al., 2007; Haynes et al., 2010). Therefore, HAF-1 appears to act as an upstream sensor that monitors the existence of mitochondrial proteotoxic stress to activate ATFS-1. In mammals, the IMM harbors three ABC transporters; ABCB7, ABCB8 and ABCB10 (Liesa et al., 2012; Schaedler et al., 2015). To date, the exact substrates of these transporters have not been determined, but the genetical ablation of these transporters in mice results in the dysregulation of intracellular iron-cofactor metabolism (Hyde et al., 2012; Ichikawa et al., 2012; Pondarre et al., 2006; Pondarre et al., 2007; Yamamoto et al., 2014).

Mitochondria play a critical role in intracellular iron metabolism, as these organelles harbor the biosynthetic pathways for two major iron-containing cofactors, heme and iron-sulfur cluster (ISC). These iron-containing cofactors that are produced at mitochondria, are ultimately incorporated into their partner proteins within the mitochondria, the cytosol or within other organelles. Therefore, it is likely that mitochondria communicate with these other compartments to coordinately regulate cellular and mitochondrial iron metabolism. However, to date, the molecular mechanisms underlying this coordination, including mitochondrial proteins that can respond to alterations in iron levels, has not been fully described.

Here, we show that mitochondrial import of DELE1 and its subsequent protein stability are regulated by intracellular iron availability. Iron deficiency-induced mitochondrial import arrest of DELE1 enabled DELE1 to escape from the degradation by the matrix-localized protease LONP1. The full-length form of DELE1 that was stabilized on the OMM and activated the HRI-ISR pathway. Importantly, neither OMA1-mediated cleavage of DELE1 nor heme deficiency was implicated in this pathway. Instead, we demonstrated a requirement for ABCB7, an IMM-resident ABC transporter that is involved in ISC metabolism. Our findings point to a hitherto unappreciated mitochondrial-based iron-monitoring system regulated by the iron-dependent mitochondrial import of DELE1 and providing an additional layer of regulation for the HRI-mediated ISR. It also underscores the importance of mitochondrial protein import regulation as a stress-sensing mechanism.

## Results

### DELE1 is a short-lived protein that is degraded by LONP1 after mitochondrial import

Previous studies have implied that the protein stability of DELE1 might be post-translationally regulated in cultured cells (Harada et al., 2010). Currently available antibodies fail to detect endogenous DELE1 (Fessler et al., 2020; Guo et al., 2020). Therefore, to monitor endogenous DELE1 protein stability, we performed cycloheximide (CHX)-chase experiments using HEK293T cells expressing an endogenously HA-tagged DELE1 (Fessler et al., 2020). On immunoblotting, DELE1 underwent rapid degradation within minutes of CHX treatment (Fig. 1A). In contrast, expression levels of other mitochondrial proteins such as TOM40, TIM50 and TIM23 did not change during this chase period (Fig. 1A). This suggests that DELE1 is actively degraded under basal conditions. This observation is consistent with a very recent study of DELE1 (Fessler et al., 2022). To identify the steady state degradation mechanism of DELE1, we first examined the effects of the proteasome inhibitor (MG132) and the autophagy inhibitor (Bafilomycin A1). Although these inhibitors accumulated their known substrates (the cleaved form of PINK1 for the proteasome and p62 for autophagy) (Sekine and Youle, 2018; Ueno and Komatsu, 2020), they did not enhance DELE1 expression (Fig. 1B), indicating that DELE1 is likely degraded independent of the proteasome and macroautophagy. Because DELE1 is a mitochondrial protein, we speculated that mitochondrial proteases are likely involved in its basal degradation. We thus conducted a small scale screen using siRNAs for reported mitochondrial proteases including PARL, OMA1, YME1L, and HTRA2, which are inner mitochondrial membrane (IMM) or intermembrane space (IMS)-resident proteases, and LONP1, CLPP, AFG3L2, and SPG7, which are mitochondrial matrix-resident proteases (Fig. 1C and 1D) (Deshwal et al., 2020). This targeted screen found that knockdown of LONP1, the matrix-localized ATP-dependent serine protease (Szczepanowska and Trifunovic, 2021), had the most pronounced effects on DELE1 (Fig. 1D). We also observed partial restoration of DELE1 expression in cells with knockdown of AFG3L2 and SPG7, that comprise the matrix-facing m-AAA protease (Fig. 1D) (Deshwal et al., 2020). In contrast, none of siRNAs for IMM or IMS proteases affected DELE1 stability (Fig. 1C). These observations suggest that DELE1 is degraded in the mitochondrial matrix predominantly in a LONP1-dependent manner. We confirmed this by an independent siRNA that targets a distinct region of the *LONP1* mRNA. Both LONP1 siRNAs increased DELE1 at steady state, and almost completely prevented DELE1 degradation during the CHX-chase (Fig. 1E). Thus, our findings indicate that DELE1 is constitutively degraded in a LONP1-dependent manner soon after mitochondrial import.

**Figure 1.**
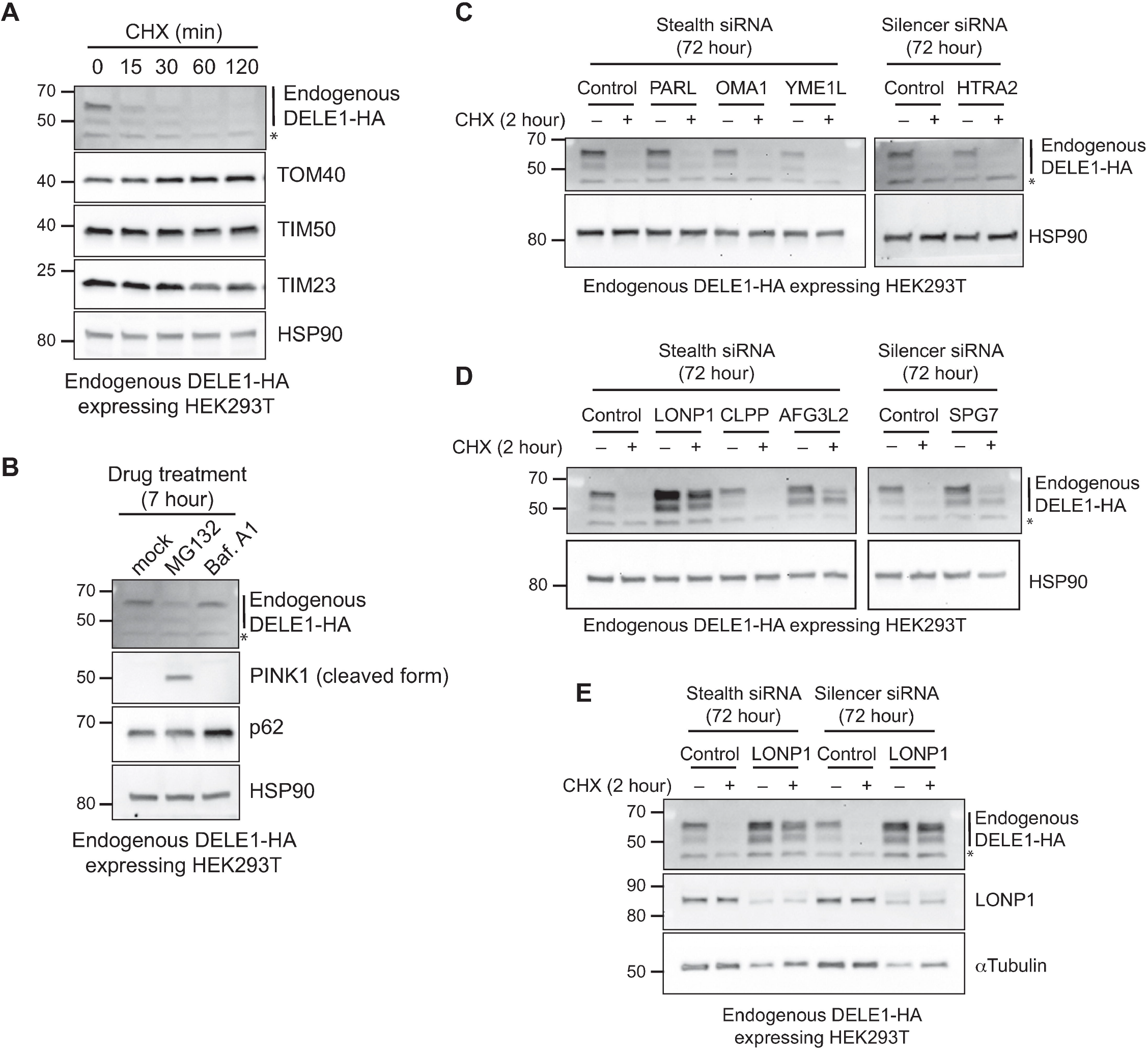
DELE1 is a short-lived protein that is degraded by LONP1 after mitochondrial import. (A) Endogenously DELE1-HA expressing HEK293T cells were treated with 10 μg/ml cycloheximide (CHX) for the indicated time periods. The lysates were analyzed by immunoblotting (IB) with the indicated antibodies. HSP90 is shown as a loading control. (B) Endogenous DELE1-HA HEK293T cells were treated with 10 μM MG132 or 200 nM Bafilomycin A1 (Baf. A1) for 7 hours. The lysates were analyzed by IB. (C-E) Endogenous DELE1-HA HEK293T cells were transfected with indicated siRNAs for 72 hours and 10 μg/ml CHX was treated for the last 2 hours before harvest. The lysates were analyzed by IB. αTubulin is shown as a loading control. *; non-specific bands.

### Iron deficiency stabilizes DELE1

LONP1 is known to function as a quality control protease that degrades misfolded and oxidatively damaged proteins in the mitochondrial matrix (Szczepanowska and Trifunovic, 2021). It has been also shown that LONP1 is actively involved in various mitochondrial processes through the regulated proteolysis of specific substrates (Szczepanowska and Trifunovic, 2021). Thus, we hypothesized that the LONP1-mediated degradation of DELE1 may be involved in some physiological response. We first tried to identify stimuli that promote DELE1 stability. Considering that DELE1 is an activator of the heme-responsive kinase HRI (Fessler et al., 2020; Guo et al., 2020), we tested heme and iron-related stimuli and found that two iron chelators, deferoxamine (DFO) and deferiprone (DFP) increased DELE1 protein levels (Fig. 2A). These compounds have been shown to decrease both cytosolic and mitochondrial iron levels (Fujimaki et al., 2019; Hara et al., 2020), and we confirmed their effects by monitoring the increase in the cytosolic iron-responsive protein IRP2 (Hentze et al., 2010) (Fig. 2A). In contrast, neither heme (hemin) nor a heme biosynthesis inhibitor (succinyl acetone, SA) affected DELE1 expression, while the heme-responsive proteins ALAS1 and HMOX1 were increased by SA and hemin, respectively (Fig. 2A). Whereas the mitochondrial uncoupler CCCP induced the cleavage of DELE1 as previously reported (Fessler et al., 2020; Guo et al., 2020) (Fig. 2A), DFO and DFP treatment appeared to preferentially promote the accumulation of a full-length form of DELE1 (Fig. 2A). Similar results were observed in a HeLa cell line that expresses a tetracycline (tet)-inducible DELE1-HA (Fig. 2B). DELE1 expression was enhanced in a DFO dose-dependent manner, similar to the response of IRP2. Furthermore, pre-treatment with DFO or DFP significantly stabilized the full-length form of DELE1 following a CHX-chase (Fig. 2C, lane 4-9). In contrast, pre-treatment with CCCP stabilized DELE1, but as the cleaved form (Fig. 2C, lane 10-12). Collectively, this implies that full-length DELE1 stability is regulated by iron availability.

**Figure 2.**
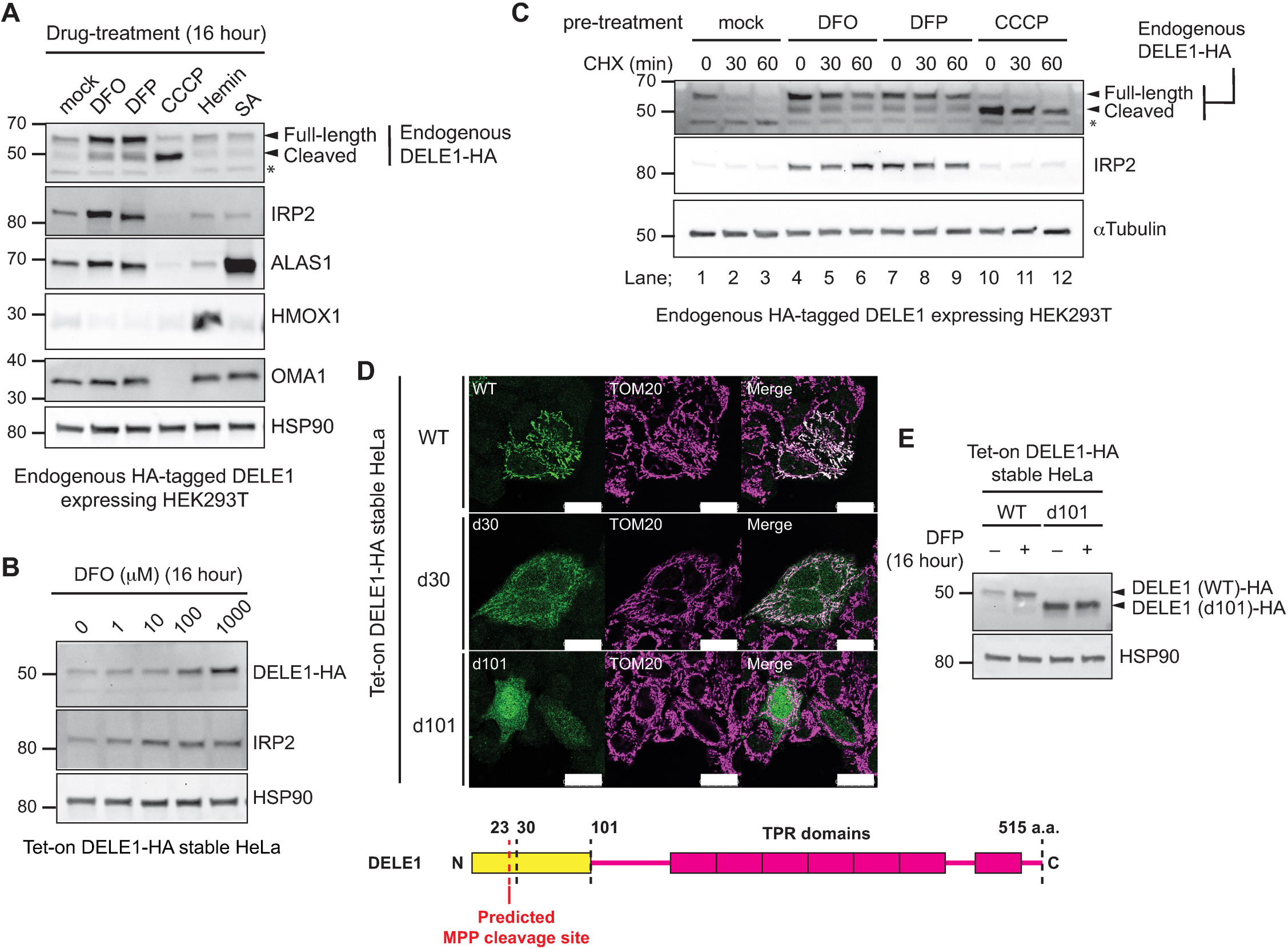
Iron deficiency stabilizes DELE1. (A) Endogenous DELE1-HA HEK293T cells were treated with 1 mM Deferoxamine (DFO), 1 mM Deferiprone (DFP), 10 μM CCCP, 20 μM Hemin or 1 mM succinyl acetone (SA) for 16 hours. The lysates were analyzed by IB with the indicated antibodies. (B) Tet-on DELE1-HA stable HeLa cells were treated with the indicated concentration of DFO for 16 hours. 1 μg/ml Doxycycline was added 8 hours prior to DFO treatment to induce the expression of DELE1-HA. The lysates were analyzed by IB. (C) Endogenous DELE1-HA HEK293T cells were pre-treated with 1 mM DFO, 1 mM DFP for 16 hours, or 10 μM CCCP for 1.5 hours and were subsequently subjected to the CHX chase with the indicated periods. The lysates were analyzed by IB. (D) Immunocytochemistry (ICC) for Tet-on DELE1 (wild-type, WT)-HA, DELE1 (delta 30, d30)-HA, DELE1 (delta 101, d101)-HA stable HeLa cells. Scale bars; 25 μm. A schematic representation of the domain structure of DELE1 is shown at the bottom. (E) Tet-on DELE1 WT-HA or DELE1 d101-HA stable HeLa cells were treated with 1 mM DFP for 16 hours. 1 μg/ml Doxycycline was added 8 hours prior to DFO treatment to induce the expression of DELE1 WT-HA or DELE1 d101-HA. The lysates were analyzed by IB.

We also examined whether the mitochondrial localization of DELE1 is required for its iron responsiveness. DELE1 has a predicted mitochondrial targeting sequence (MTS) at its N-terminus and the MTS cleavage site is predicted to be located after amino acid 23 by the web tool for MTS prediction: MitoFates (Fig. 2D) (Fukasawa et al., 2015), which has been experimentally verified (Fessler et al., 2022). We confirmed that tet-induced DELE1 wild-type (WT) is localized to mitochondria (Fig. 2D). Consistent with previous reports (Fessler et al., 2020; Guo et al., 2020), a DELE1 mutant that lacked the predicted MTS (DELE1 d30) only partially prevented mitochondrial localization of DELE1, but deletion of the N-terminal 101 amino acids of DELE1 (DELE1 d101) completely blocked mitochondrial localization (Fig. 2D). These results suggest that mitochondrial localization of DELE1 is regulated by a longer N-terminal region encompassing the predicted MTS. Unlike DELE1 WT, the DELE1 d101 mutant lost the ability to accumulate in response to iron chelation (Fig. 2E), suggesting that DELE1 responds to iron deficiency through its mitochondrial targeting N-terminus.

### DELE1 is stabilized on mitochondria by an iron deficiency-dependent mitochondrial import arrest

We next sought to gain greater understanding of the mechanisms underlying iron-deficiency-dependent DELE1 stabilization. As DELE1 was degraded by LONP1 (Fig. 1D and 1E), we first examined whether the DELE1 stabilization was mediated through inactivation of the LONP1 protease activity. Treatment of iron chelators significantly decreased mitochondrial ISC-containing proteins including SDHB and LIAS (Figure 3A, lane 1-3), suggesting that iron deprivation affected the mitochondrial ISC biosynthesis pathway. It has been reported that misfolded SDHB due to the failure of ISC insertion undergoes degradation by LONP1 (Maio et al., 2016). Indeed, LONP1 knockdown restored the expression of SDHB as well as that of LIAS in iron-chelated cells (Fig. 3A, lane 4-9), suggesting that LONP1 mediates the degradation of these ISC proteins. Thus, it appears that LONP1 is still active for degrading misfolded ISC proteins under iron deficient conditions, although DELE1 appears to selectively escape from LONP1-mediated degradation.

**Figure 3.**
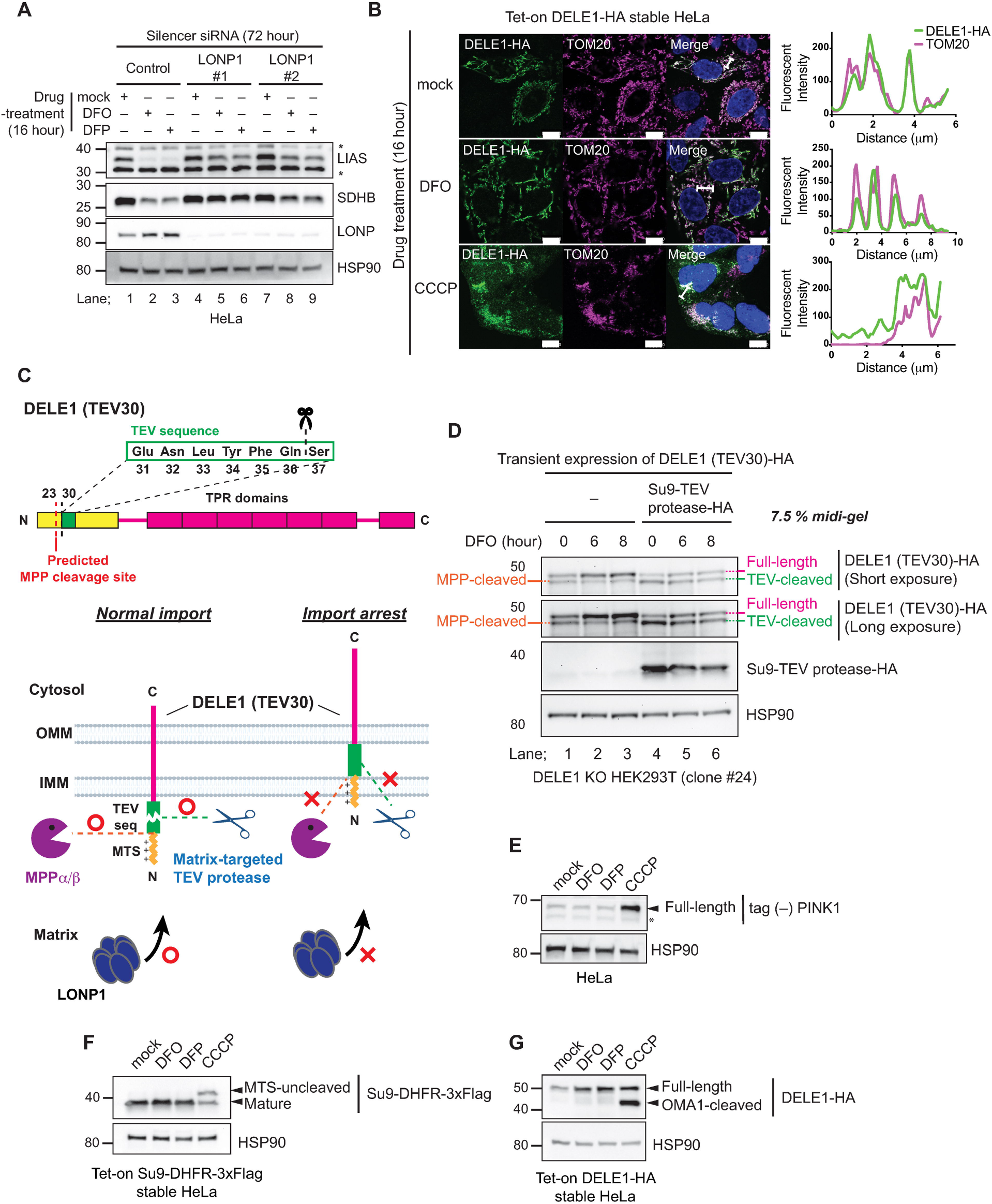
DELE1 is stabilized on mitochondria by an iron deficiency-dependent mitochondrial import arrest. (A) HeLa cells were transfected with control or LONP siRNAs for 72 hours. Cells were treated with 1 mM DFO or 1 mM DFP for the last 16 hours. The lysates were analyzed by IB with the indicated antibodies. (B) Tet-on DELE-HA stable HeLa cells were treated with 1 mM DFO or 20 μM CCCP for 16 hours. The subcellular localization of DELE1-HA was determined by immunocytochemistry (ICC). Scale bars; 10 μm. Line profiles for indicated fluorescent intensities determined along the white lines are shown to the right. (C) Schematic representation for the TEV cleavage assay. (Top) Insertion of the TEV cleavage sequence after the predicted MPP cleavage site of DELE1 [DELE1(TEV30)]. (Bottom) Suppression of the cleavage of DELE1 by import arrest. (D) DELE1 knockout (KO) cells (clone #24) were transfected with the TEV sequence-inserted DELE1 (DELE1 (TEV30)-HA) with or without the matrix-targeted TEV protease (Su9-TEV protease-HA). After 24 hours, cells were treated with 1 mM DFO for the indicated time periods. The lysates were separated in 7.5% polyacrylamide Midi gel and analyzed by IB. (E to G) HeLa cells transiently transfected with tag (–) PINK1 (E), Tet-on Su9-DHFR-3×Flag stable HeLa cells (F), or Tet-on DELE1-HA stable HeLa cells (G) were treated with 1 mM DFO, 1 mM DFP or 20 μM CCCP for 16 hours. 1 μg/ml Doxycycline was added 8 hours prior to DFO treatment to induce the expression of Su9-DHFR-3xFlag. The lysates were analyzed by IB. *; non-specific bands.

It has been reported that mitochondrial stress (e.g., CCCP treatment) induces cleavage of DELE1 by OMA1 and that cleaved DELE1 is released into the cytosol (Fessler et al., 2020; Guo et al., 2020). We confirmed this observation in our tet-on DELE1-HA HeLa cells (Fig. 3B, lower panel). In contrast, we found that DELE1 was detected predominantly on mitochondria after treatment with DFO (Fig. 3B, middle panel), indicating that DELE1 does not substantially translocate to the cytosol during iron deficiency. This is consistent with the observation that iron chelators did not induce the cleavage of DELE1 but accumulated the full-length form of DELE1 (Fig. 2A–2C). To determine the precise localization of DELE1 within mitochondria before and after iron chelation, we performed a TEV protease assay. To monitor the entrance of the N-terminus of DELE1 into the matrix, the 7 amino acids TEV sequence (that is recognized and cleaved by TEV protease) was inserted immediately after the predicted MTS of DELE1 [DELE1(TEV30)] (Fig. 3C). We transiently expressed DELE1(TEV30) with a matrix-targeted TEV protease (Su9-TEV protease; Su9 is a matrix targeting mitochondrial signal sequence of subunit 9 of the F0-ATPase of *Neurospora crassa*) in DELE1 KO HEK293T cells (Fig. S1), and subsequently treated the cells with DFO. In this setting, if the N-terminal TEV sequence of DELE1 reaches the matrix, the N-terminal 36 amino acids of DELE1 should be cleaved by the Su9-TEV protease (Fig. 3C, lower panel). In the absence of Su9-TEV protease, the N-terminal MTS of DELE1 should be cleaved by endogenous MTS-cleaving peptidase (MPP) in the matrix (Fessler et al., 2022). Indeed, DELE1(TEV30) appeared as a doublet under steady state conditions when resolved on an SDS-PAGE 7.5% acrylamide gel (Fig. S2). Upon knockdown of MPP proteases, the lower band of DELE1(TEV30) disappeared (Fig. S2), demonstrating that we could separate the full-length form and an MPP-cleaved form of DELE1(TEV30) using this approach (Fig. S2). Notably, following iron chelation with DFO, the MTS cleavage of DELE1(TEV30) was partially blocked (Fig. 3D, lane 1-3), suggesting that the N-terminus of DELE1 is inaccessible to the matrix-resident MPP under iron deficient conditions (Fig. 3C, lower panel). Consistently, cleavage of DELE(TEV30)-HA by matrix TEV was partially blocked following iron chelation with DFO (Fig. 3D, lane 4 vs. lane 5 and 6). These observations suggest that under iron deficient conditions the N-terminal region of DELE1 reaches the matrix less readily than in the steady state due to mitochondrial import arrest (Fig. 3C, lower panel). As the protease responsible for DELE1 steady-state degradation, LONP1 (Fig. 1 D and E), resides in mitochondrial matrix, this iron deficiency-induced mitochondrial import arrest of DELE1 may enable DELE1 to escape from the LONP1-mediated degradation, thereby promoting stabilization of full-length DELE1 (Fig. 2C and Fig. 3C, lower panel).

Mitochondrial membrane potential across the IMM provides a driving force for the mitochondrial import of MTS-containing proteins (Wiedemann and Pfanner, 2017). Consistently, mitochondrial depolarization by CCCP therefore induced the accumulation of the full-length form of PINK1 and Su9-DHFR as reported previously (Fig. 3E and 3F, CCCP-treated lanes) (Sekine et al., 2019). Although DELE1 is cleaved by OMA1 under CCCP-treated conditions, the total amount of DELE1 protein increased (Fig. 2A and Fig. 3G, CCCP-treated lanes). This indicates that like other mitochondrial proteins with a canonical MTS, the mitochondrial import of DELE1 depends on mitochondrial membrane potential. In contrast, while DELE1 was stabilized by iron deficiency (Fig. 2A and Fig. 3G, DFO or DFP-treated lanes), both PINK1 and Su9-DHFR were insensitive to this perturbation (Fig. 3E and 3F, DFO or DFP-treated lanes). Thus, iron chelation-dependent import arrest is not a general means of regulation for MTS-containing proteins and may be specific for DELE1.

### DELE1 activates an HRI-mediated ISR following iron deficiency

We next sought to identify the physiological role for the iron-dependent regulation of DELE1. Because previous observations have implicated DELE1 as a mediator of HRI-dependent ISR induction following mitochondrial stress (Fessler et al., 2020; Guo et al., 2020), we examined the possibility that stabilized DELE1 in response to iron deficiency also activates the HRI-ISR pathway. We first examined whether iron chelation activates the ISR. As shown in Fig. 4A, treatment with the iron chelators DFO or DFP increased phosphorylation of eIF2α and expression of ATF4 in a time dependent manner. The time course of ATF4 accumulation by iron chelation was slower than that induced by CCCP. The induction of ATF4 by DFO was concentration dependent and also evident when we employed a flow cytometric quantitative analysis using an ATF4 reporter (Fig. 4B and 4C) (Guo et al., 2020). Moreover, the increase in ATF4 expression was attenuated by ISRIB, a chemical inhibitor of ISR (Fig. S3A) (Sekine et al., 2015; Sidrauski et al., 2013), indicating that iron deficiency activates the ISR. In contrast, the heme biosynthesis inhibitor SA did not activate the ISR at least within time points investigated (Fig. 3D). Moreover, addition of hemin to the culture media did not suppress the iron chelation (and CCCP)-induced ISR activation while it did, as expected, induce HMOX1 expression (Igarashi and Sun, 2006) (Fig. 4E). These results suggest that iron deficiency can activate the ISR independent of heme deficiency.

**Figure 4.**
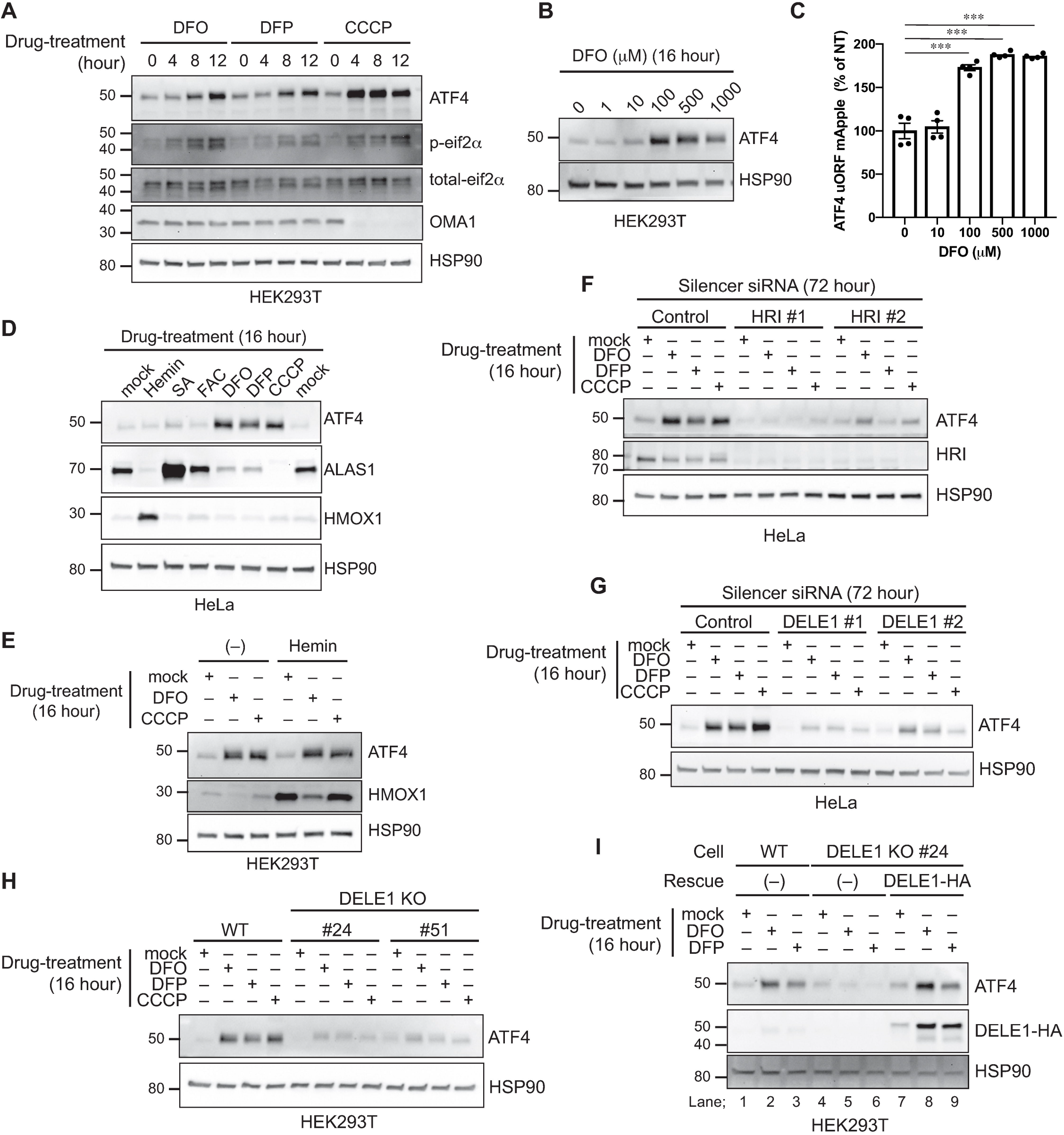
DELE1 activates an HRI-mediated ISR following iron deficiency. (A, B) HEK293T cells were treated with 1 mM DFO, 1 mM DFP or 10 μM CCCP for the indicated time periods (A) or were treated with the indicated concentrations of DFO for 16 hours (B). The lysates were analyzed by IB with the indicated antibodies. (C) ATF4 reporter (ATF4 uORF mApple)-expressing HeLa cells were treated with the indicated concentrations of DFO for 16 hours and were subjected to flow cytometry analysis. Date are shown as mean ± S.D. *** P = 0.0001 (one-way ANOVA followed by Dunnett’s multiple comparison’s test). (D) HeLa cells were treated with 20 μM Hemin, 1 mM succinyl acetone (SA), 100 μg/ml ferric ammonium citrate (FAC), 1 mM DFO, 1 mM DFP, or 20 μM CCCP for 16 hours. The lysates were analyzed by IB. (E) HEK293T cells were treated with 1 mM DFO or 10 μM CCCP for 16 hours with or without 20 μM Hemin. The lysates were analyzed by IB. (F, G) HeLa cells were transfected with siRNAs targeting HRI (F) or DELE1 (G) for 72 hours. Cells were treated with 1 mM DFO, 1 mM DFP, or 20 μM CCCP for the last 16 hours. The lysates were analyzed by IB. (H) HEK293T WT or two independent DELE1 KO cell lines (#24 and #51) were treated with 1 mM DFO, 1 mM DFP, or 10 μM CCCP for 16 hours. The lysates were analyzed by IB. (I) DELE1-HA was expressed in DELE1 KO cells through lentivirus infection for 24 hours. Cells were then treated with 1 mM DFO or 1 mM DFP for 16 hours. The lysates were analyzed by IB.

We next tested the involvement of HRI and DELE1 in iron chelation-induced ISR activation. Knockdown experiments revealed that both HRI and DELE1 were required for DFO and DFP-induced ISR activation (Fig. 4F and 4G). The knockdown efficiency of DELE1 was confirmed by RT-PCR (Fig. S3B). We further confirmed this observation in our DELE1 KO cell lines (Fig. S1). Genetic ablation of DELE1 suppressed iron chelation-induced ISR activation (Fig. 4H and 4I, lane 1-6). Furthermore, exogenous expression of DELE1 rescued the ISR activation in DELE1 KO cells (Fig. 4I, lane 7-9). Thus, DELE1is required for activation of the HRI-ISR pathway in iron-deficient cells.

### DELE1 on mitochondrial surface activates HRI

Mitochondrial depolarization caused by CCCP induces cleavage of DELE1 by OMA1, which promotes the cytoplasmic translocation of DELE1, thereby enabling DELE1 to interact with HRI in the cytosol (Fessler et al., 2020; Guo et al., 2020). However, we observed that iron chelators led to the accumulation of the full-length form of DELE1 (Fig. 2A-C), and DELE1 localized to the mitochondria during iron deficiency (Fig. 3B). These observations suggest that iron chelators activate the DELE1-HRI pathway in a mechanism distinct from CCCP. Indeed, unlike CCCP, neither DFO nor DFP induced mitochondrial depolarization (Fig. 5A). OMA1 is a stress-responsive protease whose proteolytic activity is enhanced by mitochondrial stress-inducing agents including CCCP (Ehses et al., 2009; Head et al., 2009). Stress-dependent OMA1 activation occurs through self-cleavage that eventually leads to the degradation of OMA1 (Baker et al., 2014; Zhang et al., 2014). Consistent with the absence of mitochondrial depolarization (Fig. 5A), iron chelators appeared not to induce OMA1 activation, which was monitored by processing of OMA1 itself, as well as its known substrates OPA1 and PGAM5 (Ehses et al., 2009; Head et al., 2009; Sekine et al., 2012; Wai et al., 2016) (Fig. 5B). Moreover, knockdown of OMA1 did not suppress iron chelation-induced ISR activation (Fig. 5C), indicating that OMA1 is not required for iron-deficiency-induced HRI activation. These results suggest that in iron-depleted conditions, DELE1 activates HRI by a distinct mechanism from how HRI is activated during mitochondrial stress.

**Figure 5.**
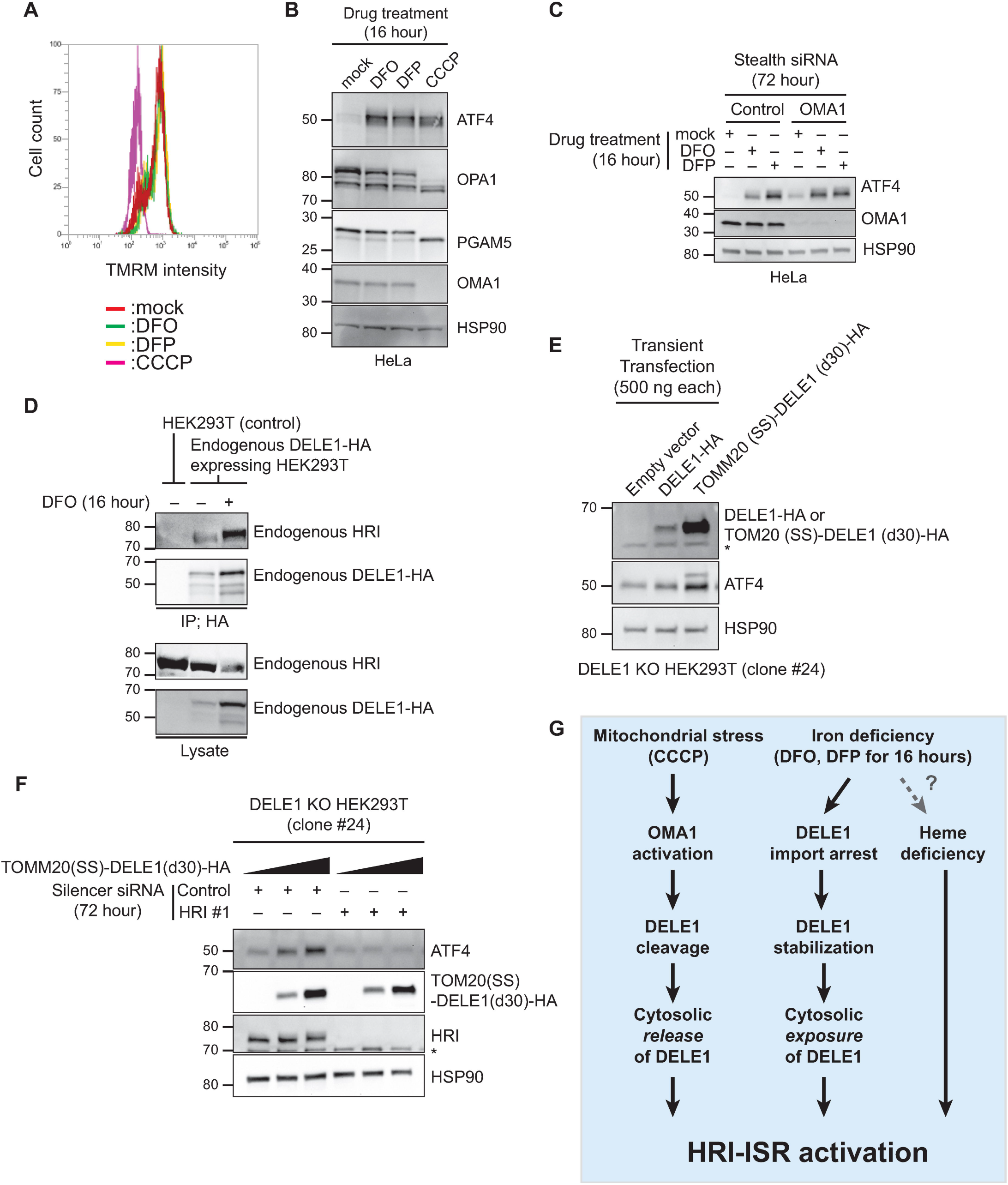
DELE1 on mitochondrial surface activates HRI. (A) HeLa cells were treated with 1 mM DFO, 1 mM DFP or 20 μM CCCP for 16 hours, and with 20 nM TMRM for the last 15 min. Cells were subjected to flow cytometry analysis to determine mitochondrial membrane potential. (B) HeLa cells were treated with 1 mM DFO, 1 mM DFP or 20 μM CCCP for 16 hours. The lysates were analyzed by IB with the indicated antibodies. (C) HeLa cells were transfected with siRNAs for control or OMA1 for 72 hours, and were treated with 1 mM DFO or 1 mM DFP for the last 16 hours. The lysates were analyzed by IB. (D) Endogenous DELE1-HA was immunoprecipitated with anti-HA antibody from the indicated HEK293T cells with or without 16 hours DFO treatment. The samples were subjected to IB. IP; immunoprecipitation. (E) DELE1 KO HEK293T (clone #24) cells were transiently transfected with DELE1-HA or the OMM-tethered DELE1 [TOMM20 (SS)-DELE1 (d30)-HA]. 500 ng of each plasmid was transfected for 24 hours. The lysates were analyzed by IB. (F) DELE1 KO HEK293T (clone #24) cells were transfected with siRNAs for control or HRI for 72 hours. Cells were then transiently transfected with 0, 500, or 1000 ng of the TOMM20 (SS)-DELE1 (d30)-HA-expressing plasmid for the last 24 hours. The lysates were analyzed by IB. (G) Multiple pathways to activate HRI-mediated ISR. See text for details. *; non-specific bands.

The C-terminal region of DELE1 contains 7 repeats of the TPR domain, known to be involved in protein-protein interactions (Fig. 2D). Previous studies have revealed that cytosolic expression of the TPR domain region of DELE1 is sufficient for the activation of HRI-ISR pathway (Fessler et al., 2020; Guo et al., 2020). Moreover, cytosolic full length DELE1 has recently been shown to be able to bind and activate HRI (Fessler et al., 2022). Our immunoprecipitation assay revealed that with the increase in DELE1 upon iron chelation, HRI bound to DELE1 also increased (Fig. 5D). We hypothesized that the mitochondrial import arrest of DELE1 resulted in retention of the C-terminal TPR domains on mitochondrial surface (Fig. 3C, lower panel), allowing for folding of the TPR domains and interaction with cytosolic HRI. We thus wanted to test if DELE1 accumulation on the mitochondrial surface, too, is sufficient for HRI activation. To this end, we tethered DELE1 to the OMM by fusing DELE1(d30) with a signal sequence (SS) of the OMM-localized protein TOM20, and examined whether this fusion protein can activate the HRI-ISR pathway. The OMM-tethered DELE1 [TOMM20(SS)-DELE1(d30)] was more stable than DELE1 WT (Fig. 5E), which is consistent with our observation that DELE1 is degraded in the matrix (Fig. 1D and 1E). Under these conditions, the OMM-tethered DELE1 induced ATF4 expression without iron chelation (Fig. 5E), suggesting that the OMM-tethered DELE1 acts as a constitutively active form for ISR induction. Furthermore, knockdown of HRI abrogated this ISR activation (Fig. 5F). These results suggest that stabilization of DELE1 on the OMM can trigger the HRI-ISR pathway in iron deficiency. This pathway does not require the OMA1-mediated DELE1 cleavage and seems to be independent from the previously suggested heme deficiency-induced activation of HRI (Fig. 5G). A very recent study uncovered that mitochondrial import stress (e.g., knockdown of mitochondrial import machineries or overexpression of mitochondrial precursor proteins) activates DELE1-HRI-ISR pathway (Fessler et al., 2022). Of note, ISR activation in this context also does not require OMA1, and it is mediated by full-length DELE1 accumulated in the cytosol (Fessler et al., 2022). Therefore, DELE1 appears to activate the HRI-ISR pathway without OMA1-mediated cleavage under certain conditions, when its C-terminal TPR domain has a physical access to cytosolic HRI.

### IMM-resident ABC transporter ABCB7 is required for the iron deficiency-induced ISR activation

Considering that DELE1 import appears to be uniquely regulated when compared to other MTS-containing proteins (Fig. 3E-3G), we next sought to identify specific factors that are required for the iron deficiency-induced import arrest of DELE1 and subsequent activation of DELE1-HRI-ISR pathway. Recent studies have revealed that LONP1 physically and functionally associates with mitochondrial import machineries (Matsushima et al., 2021; Shin et al., 2021). Thus, we first examined the possibility that LONP1 is involved in the iron deficiency-induced ISR activation through regulating the DELE1 import status. However, we did not see any defects in iron deficiency-induced ISR activation following LONP1 knockdown (Fig. 6A). As previously reported (Fessler et al., 2020), the LONP1 knockdown slightly activated the ISR at steady state (Fig. 6A), which may imply the presence of mitochondrial proteotoxic stress in LONP1-deficient cells, as LONP1 is a key quality control protease (Szczepanowska and Trifunovic, 2021).

**Figure 6.**
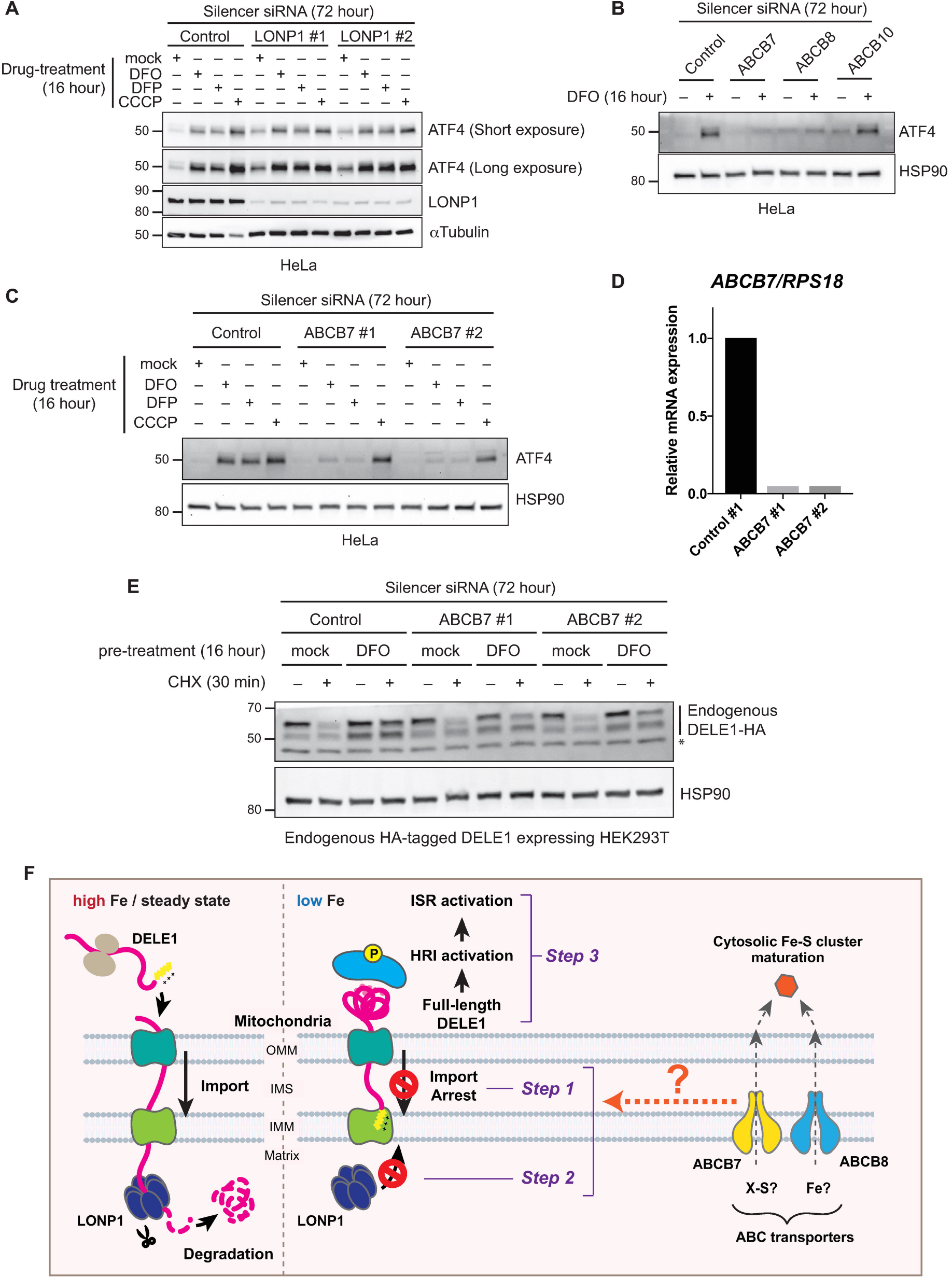
IMM-resident ABC transporter ABCB7 is required for the iron deficiency-induced ISR activation. (A) HeLa cells were transfected with the indicated siRNAs for 72 hours. Cells were treated with 1 mM DFO, 1 mM DFP, or 20 μM CCCP for the last 16 hours. The lysates were analyzed by IB with the indicated antibodies. (B) HeLa cells were transfected with the indicated siRNAs for 72 hours. Cells were treated with 1 mM DFO for the last 16 hours. The lysates were analyzed by IB. (C) HeLa cells were transfected with the indicated siRNAs for 72 hours. Cells were treated with 1 mM DFO, 1 mM DFP, or 20 μM CCCP for the last 16 hours. The lysates were analyzed by IB. (D) Knockdown efficiency of ABCB7 in (C). (E) Endogenous DELE1-HA HEK293T cells were transfected with the indicated siRNAs for 72 hours. Cells were pre-treated with 1 mM DFO for 16 hours, followed by the CHX chase for 30 min. The lysates were analyzed by IB. *; non-specific bands. (F) A proposed model of the iron-deficiency-induced mitochondrial import regulation of DELE1 and the subsequent activation of the HRI-ISR pathway. See text for details.

In *C. elegans*, the mitochondrial import regulation of ATFS-1, a unique transcription factor that harbors both MTS and nuclear localization signal (NLS), plays a critical role in mitochondrial-nuclear communication under mitochondrial proteotoxic stress (mtUPR) (Nargund et al., 2012). It has been suggested that the mitochondrial stress-activated ISR is a functional counterpart of mtUPR in mammalian cells (Eckl et al., 2021; Fiorese et al., 2016; Quiros et al., 2017). ATFS-1 is constitutively degraded by LONP-1 (Nargund et al., 2012), the *C. elegans* ortholog of mammalian LONP1 that we identified for the steady state degradation of DELE1 (Fig. 1D and 1E). These observations point to a mechanistic similarity between mtUPR and the DELE1-HRI-ISR pathway. HAF-1, the IMM-resident ABC transporter, was identified as a critical factor for the mitochondrial import arrest of ATFS-1 (Haynes et al., 2010; Nargund et al., 2012). Although the precise molecular mechanism still remains elusive, it has been suggested that HAF-1 exports peptides derived from unfolded proteins that are digested by the matrix resident protease CLPP, which ultimately prevents the mitochondrial import of ATFS-1 (Haynes et al., 2007; Haynes et al., 2010; Nargund et al., 2012). In mammals, there are three ABC transporters (ABCB7, ABCB8 and ABCB10) in the IMM (Liesa et al., 2012; Schaedler et al., 2015). Intriguingly, all of these transporters are reported to play critical roles in the intracellular iron-cofactor metabolism. Ablation of ABCB7 or ABCB8 in mice disrupts cytosolic ISC biogenesis (Ichikawa et al., 2012; Pondarre et al., 2006), while ablation of ABCB10 in mice disrupts heme biosynthesis (Hyde et al., 2012; Yamamoto et al., 2014). Loss-of-function mutations in ABCB7 cause X-linked sideroblastic anemia with ataxia (Shimada et al., 1998), and ABCB7 was also proposed to be involved in heme biosynthesis (Maio et al., 2019; Pondarre et al., 2007). It has been suggested that ABCB7 exports a yet-unknown, sulfur-containing compound (typically designated “X-S”) from the mitochondria to the cytosol (Lill and Freibert, 2020), and ABCB8 exports iron directly or indirectly from mitochondria to the cytosol (Ichikawa et al., 2012).

Due to the similarity between the ATFS-1-mediated mtUPR response and the DELE1-mediated ISR response, along with our observation that DELE1 is an iron-responsive molecule, we hypothesized that one or more of the IMM ABC transporters may be involved in the iron depletion-induced ISR activation. As shown in Fig. 6B, the ablation of ABCB7 and ABCB8, but not ABCB10, attenuated iron deficiency-induced ISR activation. Because ABCB8 knockdown severely impaired cell growth, we focused our subsequent efforts on ABCB7. We confirmed the requirement of ABCB7 in the iron deficiency-induced ISR activation by another ABCB7 siRNA that targets a different region of *ABCB7* mRNA (Fig. 6C). We confirmed the knockdown efficiency of ABCB7 by RT-PCR (Fig. 6D). Notably, ABCB7 was selectively required for the ISR activation by iron chelators DFO and DFP but not for that induced by CCCP (Fig. 6C). Furthermore, in a CHX-chase experiment, we found that stabilization of DELE1 in response to iron chelation was less efficient in ABCB7 knockdown cells (Fig. 6E). Considering that DELE1 stabilization is presumably induced by DELE1 import arrest (Fig. 5G), this observation suggests that the iron deficiency-induced import arrest of DELE1 may require ABCB7 (and ABCB8) and also implies the possible involvement of ISC-related molecules that may be exported through these ABC transporters.

## Discussion

Based on these findings, we propose a following model of DELE1-mediated HRI activation under low iron conditions (Fig. 6F). Under steady state conditions where enough iron is available, DELE1 is constitutively degraded by the matrix-resident protease LONP1 soon after mitochondrial import, thereby maintaining low steady-state levels of DELE1 expression (Fig. 6F, left). When intracellular iron availability is limited, the mitochondrial import of DELE1 is arrested. This arrest likely requires the activity of components involved in the ISC biosynthesis pathway including the IMM-resident ABC transporters ABCB7 and ABCB8 (Fig. 6F, right, step 1). This iron deficiency-induced mitochondrial import arrest of DELE1 enables DELE1 to escape from LONP1-mediated degradation (Fig. 6F, right, step 2). As a consequence, full-length DELE1 is stabilized on the OMM, exposing its C-terminal domain to the cytosol, and thereby allowing the TPR domains to fold and interact with HRI (Fig. 6F, right, step 3).

Recent studies revealed the importance of mitochondrial import regulation in sensing various mitochondrial perturbations and triggering the appropriate stress response. These include the PINK1/Parkin-mediated mitophagy pathway triggered by mitochondrial depolarization-induced PINK1 import arrest (Sekine and Youle, 2018), and the ATFS-1-mediated mtUPR in *C. elegans*, triggered by mitochondrial proteotoxic stress-induced ATFS-1 import arrest (Anderson and Haynes, 2020). Notably, expression of both proteins is kept at a low level in the absence of stress due to their constitutive degradation. In both cases, the stress-dependent import arrest enables them to escape from this degradation to elicit a stress response. The constitutive degradation of these proteins is an energy-consuming process but allows a rapid response to stress sensed within mitochondria. Similar to PINK1 and ATFS-1, DELE1 import arrest underscores this general mechanism of mitochondrial stress sensing, in this case to sense mitochondrial iron deficiency and activate the ISR. Particularly, our findings demonstrated the mechanistic similarity between the ATFS-1-dependent mtUPR and the DELE1-dependent ISR: namely, the requirement of LONP1 for steady state degradation of the stress sensor and ABC transporters for the stress-dependent activation.

There are few examples of proteins whose mitochondrial import is regulated by iron. In our study, iron deficiency-dependent import arrest was observed only for DELE1, and not two other MTS-containing proteins we tested (Fig. 3E-3G). A recent report identified that FTMT, a mitochondrial ferritin, may also be subject to import arrest in iron deficiency, and that this regulation may be related to its role in iron depletion-induced mitophagy (Hara et al., 2020). Moreover, it has been suggested that ATFS-1 is also activated by several stresses including *P. aeruginosa* infection, in addition to the mitochondrial proteotoxic stress (Pellegrino et al., 2014). Intriguingly, the exposure of worms to *P. aeruginosa* strains lacking siderophore, a bacterial toxic compound that has an iron-chelating activity, resulted in less mtUPR activation than that induced by wild-type *P. aeruginosa* (Pellegrino et al., 2014). Thus, it is tempting to speculate that ATFS-1 import might also be sensitive to bacterial toxin-mediated iron chelation, and possibly to other iron-deficient conditions.

It has been revealed that mitochondrial localization of DELE1 relies on relatively long N-terminal region (1-100 amino acids) (Fessler et al., 2020; Guo et al., 2020) that contains a short MTS (1-23 amino acids) (Fig. 2D) (Fessler et al., 2022). We also found that the deletion of N-terminal 100 amino acids of DELE1 abrogated the response of DELE1 to iron depletion (Fig. 2E). The MTSs of PINK1 and ATFS-1 have unique properties that are not observed in other typical MTS-containing proteins, which enables their stress-dependent import regulation. Similar to DELE1, the deletion of the predicted MTS (1-34 amino acids) of PINK1 does not prevent its mitochondrial localization. Rather, it stabilizes PINK1 on the OMM through a unique second MTS that is found around 70-100 amino acids of PINK1 (Okatsu et al., 2015). In addition to these two MTSs, several critical motifs for the import regulation were reported in the N-terminal 1-150 amino acids of PINK1 (Sekine, 2020; Sekine et al., 2019). In the case of ATFS-1, the MitoFates algorithm (Fukasawa et al., 2015) predicts a relatively low likelihood of mitochondrial targeting, suggesting that the MTS of ATFS-1 is comparatively weak (Melber and Haynes, 2018). Recent studies revealed that this weak MTS of ATFS-1 is critical to its stress sensitivity (Rolland et al., 2019; Shpilka et al., 2021). In contrast to ATFS-1, the MTS prediction for DELE1 by MitoFates is relatively strong (Probability of presequence: DELE1, 0.753; ATFS-1, 0.118; PINK1, 0.996 and ATP5A, 1.0), suggesting the existence of other regulatory mechanisms. The detailed examination of the N-terminal region of DELE1 might provide a clue to delineate the iron-dependent import regulation of this protein. In this perspective, it is worth notifying that very recent detailed analyses of DELE1 import identified a stop-transfer like motif in the end of the extended MTS region (80-106 amino acids) of DELE1 (Fessler et al., 2022). This motif, which is a short hydrophobic amino acid sequence that appears to form an α-helical configuration, may slow the DELE1 import across the IMM and thereby assist the OMA1-mediated cleavage in stressed mitochondria (Fessler et al., 2022). As mentioned above, a similar motif in the extended MTS is required for the retention of PINK1 in the OMM in response to mitochondrial depolarization (Okatsu et al., 2015). Therefore, it would be interesting to test whether the iron deficiency-induced DELE1 import arrest also relies on this unique motif in the extended MTS.

We observed that the ablation of ABCB7 and ABCB8, components involved in ISC metabolism, abolished iron deficiency-induced ISR activation (Fig. 6B). Specifically, the knockdown of ABCB7 attenuated iron deficiency-induced DELE1 stabilization (Fig. 6E). Therefore, it is tempting to speculate that some ISC-related molecules, such as sulfur-containing factors and iron, exported by ABCB7 and ABCB8 are required for the mitochondrial import arrest of DELE1, thereby promoting its stability. In this respect, it is noteworthy that other small biological molecules have been reported to regulate mitochondrial import. For example, heme directly binds the MTS of ALAS, the rate limiting enzyme for heme biosynthesis, preventing its mitochondrial import, as a negative feedback mechanism to maintain appropriate intracellular heme levels (Lathrop and Timko, 1993). It will be interesting to test whether a similar direct mechanism of DELE1 import regulation occurs following iron depletion, perhaps by direct binding of an ISC-related molecule to the MTS of DELE1. Alternatively, it has been reported that the ablation of ABCB7 and ABCB8 results in the mitochondrial iron overload (Cavadini et al., 2007; Ichikawa et al., 2012; Maio et al., 2019). Therefore, we cannot rule out the possibility that the extent of iron chelation in mitochondria is less efficient in ABCB7- or ABCB8-deficient cells than in WT cells. Nevertheless, either scenario supports our conclusion that DELE1 responds to changes in intracellular iron metabolism.

In this study, we described a novel mechanism of HRI activation in response to iron deficiency that requires DELE1. The core finding of this study is the identification of DELE1 as a unique protein whose mitochondrial import and subsequent proteolysis is regulated by intracellular iron availability. The iron-dependent mitochondrial import regulation of DELE1 and its critical role in the HRI activation represent a previously unrecognized mitochondrial-based iron sensing mechanism connecting mitochondria to the cytosolic ISR. Mitochondria are the major iron containing subcellular compartment, as they harbor the biosynthetic pathways for two major iron-containing cofactors, heme and ISC. Therefore, it is not surprising that mitochondria would have their own iron monitoring system, in addition to the well-characterized iron homeostasis maintenance mechanism governed by cytosolic IRP1/2 (Hentze et al., 2010). The pro-survival roles of HRI in the erythroid lineage are well established (Han et al., 2001). As such, in erythroid cells, further analyses of the DELE1-HRI pathway may yield new therapeutic targets for various red blood cell disorders including iron-related anemias and inherited hemoglobinopathies.

## Acknowledgements

We thank Dr. Toren Finkel for critical reading our manuscript, and Dr. Richard J Youle for sharing reagents. This work was supported by University of Pittsburgh, Aging Institute Startup Seed (Y.S. and S.S.), and was supported in part by the NINDS intramural program (D.P.N.).

## Author contributions

S.S. led the project. Y.S. and S.S. wrote an initial draft of the manuscript. Y.S., R.H., and S.S. designed and performed experiments. E.F. and L.T.J. supported the project by providing endogenously HA-tagged DELE1 expressing HEK293T cells and suggestions. D.P.N. performed the flow cytometric analysis of the ATF4 reporter cell line and provided suggestions. All the authors edited the manuscript.

## Figure legends

**Supplemental Figure 1**

**Generation of DELE1 knockout (KO) HEK293T cell lines.**

A genotyping PCR result for DELE1 KO clones (Top) and the design of sgRNAs for genetic deletion of DELE1 in HEK293T cells by CRISPR-mediated genome editing (Bottom). Primers used for genotyping PCR that amplify 1190 bp in *WT DELE1* gene are also indicated (Bottom).

**Supplemental Figure 2**

**Supplemental information related to Figure 3.**

DELE1 KO HEK293T cells (clone #24) were transfected with the indicated siRNAs for 72 hours. Cells were then transiently transfected with the DELE1 (TEV30)-HA plasmid for the last 24 hours. The lysate was analyzed by SDS-PAGE. The lysates were analyzed by IB with the indicated antibodies.

**Supplemental Figure 3**

**Supplemental information related to Figure 4.**

(A) HeLa cells were treated with 1 mM DFO, 1 mM DFP or 20 μM CCCP for 16 hours in the presence or absence of the indicated concentrations of ISRIB. The lysates were analyzed by IB with the indicated antibodies. (B) HeLa cells were transfected with the indicated siRNAs for 72 hours and cDNA pools were prepared from isolated total RNAs of these cells. Quantitative PCR analysis was performed for confirming DELE1 knockdown efficiency.

**Table 1**

**Key Resource Table**

## Contact for Reagent and Resource Sharing

Further information and requests for reagents will be fulfilled by Lead Contact Shiori Sekine.

## Materials and methods

### Cell culture, transfection and reagents

HEK293T and HeLa cells were cultured in DMEM (Gibco) supplemented with 10% FBS (VWR Life Science), 10 mM HEPES (Gibco), 1 mM Sodium pyruvate (Gibco), non-essential amino acids (Gibco) and GlutaMAX (Gibco). For RNA interference, 20 nM Stealth siRNAs (Thermo Fisher Scientific) or 5 nM Silencer select siRNAs were transfected using Lipofectamine RNAi max transfection reagent (Thermo Fisher Scientific) at the same time as cell seeding. For transient transfection of plasmids, X-tremeGENE 9 DNA Transfection Reagent (Sigma-Aldrich) was used. All the siRNAs and reagents used in this study are described in Table1.

### Plasmids

All plasmids construction was performed by PCR amplification (CloneAmp HiFi PCR Premix) using appropriate primers followed by Gibson assembly (In-Fusion HD Cloning system, Clontech) into the EcoRI site of pLVX-puro vector (Clontech) or into the BamHI/NotI site of pRetroX-Tight-puro vector (Clontech). The original vector of C-terminally HA-tagged Su9-TEV protease was gifted from Dr. Thomas Langer (Max Planck Institute) (Baker et al., 2014), and sub-cloned into the HindIII-XhoI site of pcDNA3.1 (+) (Sekine et al., 2019). Mammalian Tag (–) PINK1 expression vector was kindly gifted from Dr. Noriyuki Matsuda (Tokyo Metropolitan Institute of Medical Science, Current affiliation: Tokyo Medical and Dental University) (Okatsu et al., 2012). All the plasmids used in this study are described in Table 1.

### Generation of cell lines

The generation of the endogenously HA-tagged DELE1 expressing HEK293T cells was described previously (Fessler et al., 2020). To generate stably transfected cell lines, lentiviruses (for plasmids within pLVX-puro vectors) and retroviruses (for plasmids within pRetroX-Tight-puro vector) were packaged in HEK293T cells. HeLa cells were transduced with viruses with 10 μg/ml polybrene (Sigma) then optimized for protein expression via antibiotics selection. For generating the Tet-on DELE1-HA stable HeLa cell line (Houston et al., 2021) or Tet-on Su9-DHFR-3xFlag stable HeLa cell line (Sekine et al., 2019), Retro-X-Tet-on Inducible Expression System (Clontech) was used according to the manufacturer’s instruction. For generating the ATF4 reporter cell line (ATF4_uORF_mApple-stable HeLa cells), HeLa cells were transduced with a previously reported lentiviral ISR reporter pXG237 containing two ATF4 upstream open reading frames upstream of mApple (Guo et al., 2020). Plasmid pXG237 was a gift from Dr. Martin Kampmann (University of California) (Addgene plasmid # 141281). A clonal cell line was produced by single-cell sorting into a 96-well plate. DELE1 KO HEK293T cell lines were generated using lentiCRISPRv2 system (Sanjana et al., 2014; Shalem et al., 2014). CRISPR target sites were chosen from 5’ and 3’ intron region of Exon2 of *DELE1* gene, and two sgRNAs were designed for each region; 5’ intron region (5’ - ggagaccagcagaatcacat-3’), 3’ intron region (5’ - ctcatttcctcccctagtca-3’). After the infection of lentiviruses that express hSpCas9 and DELE1 sgRNAs, infected cells were selected via the treatment with 500 μg/ml puromycin (Sigma) for 24 hours. The selected cells were subjected to single colony isolation in 96-well plates. Genomic DNA of the DELE1 KO clones (#24 and #51) used in this study were extracted with the Genotyping Buffer [100 mM Tris-HCl pH 8.0, 5 mM EDTA, 200 mM NaCl, 0.1% SDS, 0.2 mg/ml proteinase K (Thermo Fisher Scientific)] and PCR amplified using a pair of primers (forward, 5’-gtccaatggcaggagatggt-3’; reverse, 5’-aggctatcaagtaggggcaaag-3’) to confirm the deletion. All the cell lines used in this study are described in Table 1.

### Immunoblotting (IB) and Immunocytochemistry (ICC)

The procedures for IB and ICC were described previously (Houston et al., 2021). All the antibodies used in this study are described in Table 1.

### Immunoprecipitation (IP)

For immunoprecipitation, cells were lysed with 1% Triton Buffer [1% Triton-X100, 150 mM NaCl, 50 mM Tris-HCl pH 7.4, 1 mM EDTA, Phosphatase inhibitors (PhosSTOP, Sigma) and protease inhibitors (cOmplete, Sigma)]. After centrifugation, the lysate was incubated with anti-HA Magnetic beads (Pierce) for 20 min. After washing beads with 1% Triton Buffer for 4 times, immunoprecipitants were eluted from the beads by boiling with 1× NuPAGE LDS sample Buffer (Thermo Fisher Scientific) supplemented with 100 mM Dithiothreitol (DTT) (Sigma) for 3 min.

### RNA Isolation and Real-Time PCR

Procedures for RNA isolation and subsequent real-time PCR were described previously (Houston et al., 2021). All expression levels were normalized to that of RPS18 mRNA. The following RT-PCR primers were used; RPS18, (forward) 5’-cttccacaggaggcctacac-3’ and (reverse) 5’-cgcaaaatatgctggaacttt-3’, and DELE1, (forward) 5’-cccactggaaaggagtgttg-3’ and (reverse) 5’-acccacaggctccctctt-3’, and ABCB7, (forward) 5’-ccacacagacccaaaagaag-3’ and (reverse) 5’-caccacccaaaaatcccag-3’.

### Flow cytometry

To determine mitochondrial membrane potential, 20 nM TMRM (Thermo Fisher Scientific) directly added to cell culture media and incubated for 15 min. Cells were washed and replaced with normal medium followed by flow cytometry analysis using Attune NxT Acoustic Focusing Cytometer (Thermo Fisher Scientific). For the flow cytometry analysis of the ATF4 reporter cell line (ATF4_uORF_mApple-stable HeLa cells), mApple was measured on an Amnis CellStream flow cytometer (Luminex), using a 561 nm laser for excitation and a 611/31 bandwidth filter for emission. Data was analyzed from the fcs files using a custom Python 3 script and the FlowCytometryTools library.

