## Supplementary figures and images for "A mitochondrial iron-sensing pathway regulated by DELE1"

### Supplemental Figure 1

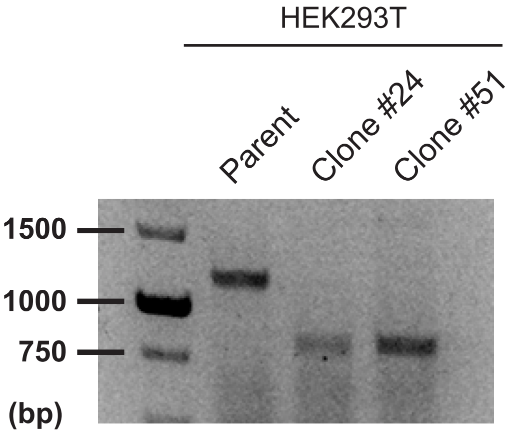


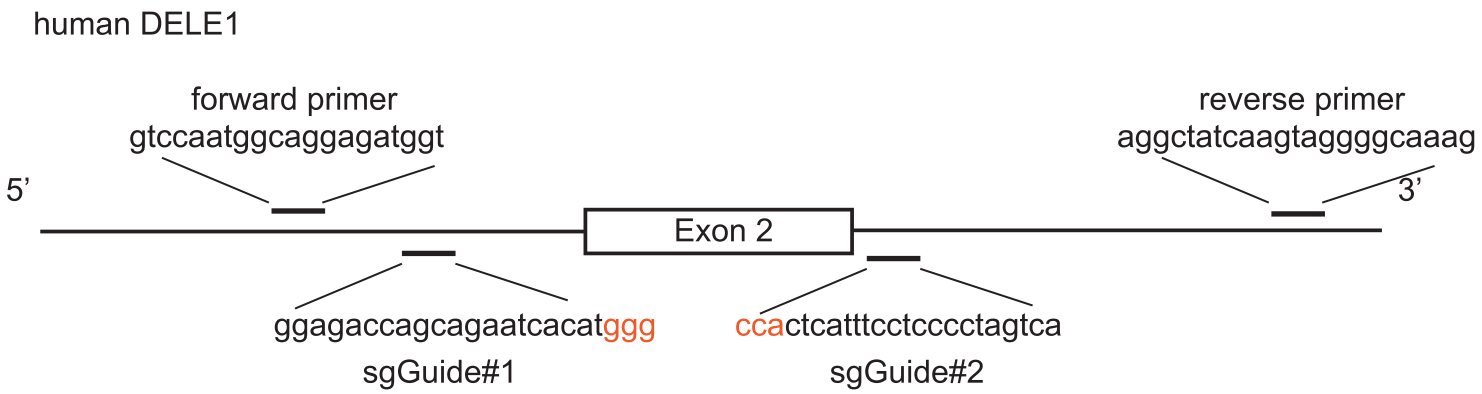


**Supplemental Figure 1**
**Generation of DELE1 knockout (KO) HEK293T cell lines.**

### Supplemental Figure 2

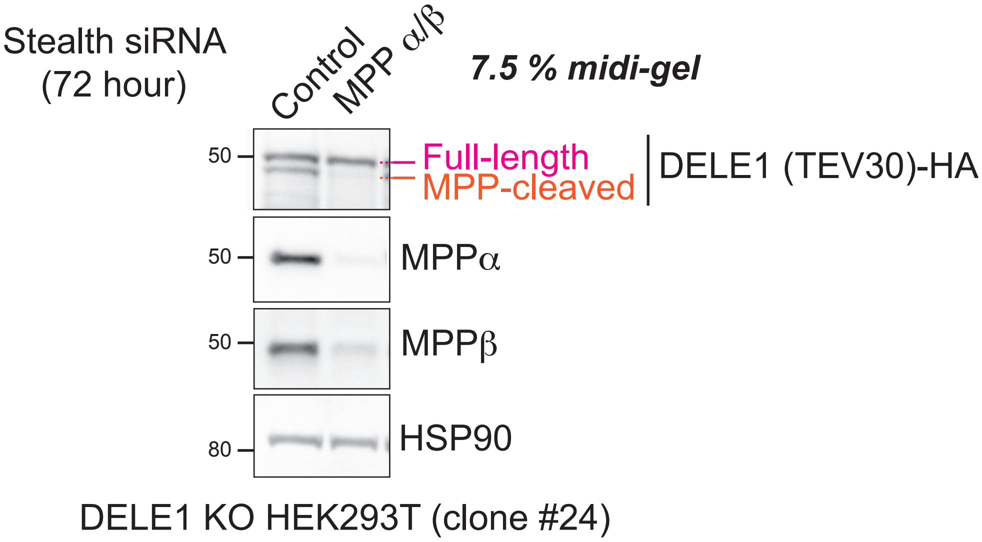


**Supplemental Figure 2**
**Suplemental data related to Fig. 3.**

### Supplemental Figure 3

A

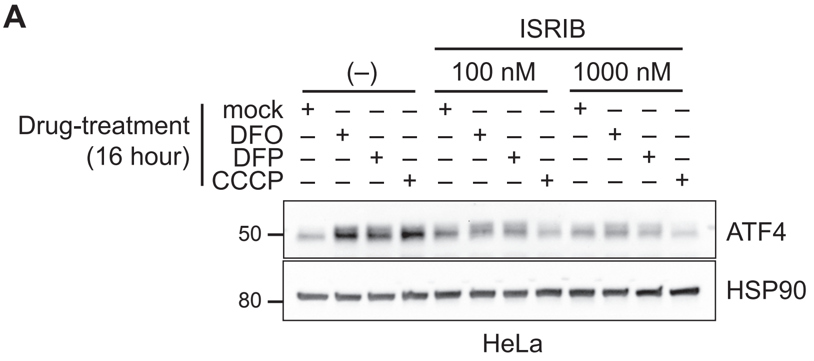


B

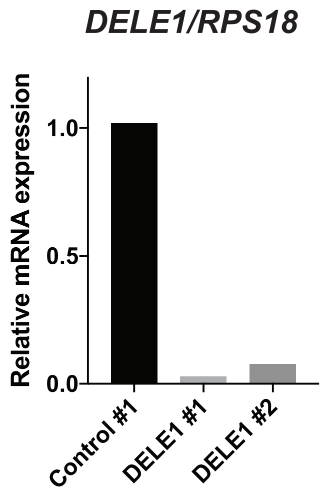


**Supplemental Figure 3**
**Supplemental data related to Figure 4.**
